# A Molecular Communication model for cellular metabolism

**DOI:** 10.1101/2023.06.28.546976

**Authors:** Zahmeeth Sakkaff, Andrew P. Freiburger, Jennie L. Catlett, Mikaela Cashman, Aditya Immaneni, Nicole R. Buan, Myra B. Cohen, Christopher Henry, Massimiliano Pierobon

## Abstract

Understanding cellular engagement with its environment is essential to control and monitor metabolism. Molecular Communication theory (MC) offers a computational means to identify environmental perturbations that direct or signify cellular behaviors by quantifying the information about a molecular environment that is transmitted through a metabolic system. We developed an model that integrates conventional flux balance analysis metabolic modeling (FBA) and MC to mechanistically expand the scope of MC, and thereby uniquely blends mechanistic biology and information theory to understand how substrate consumption is captured reaction activity, metabolite excretion, and biomass growth. This is enabled by defining several channels through which environmental information transmits in a metabolic network. The information flow in bits that is calculated through this workflow further determines the maximal metabolic effect of environmental perturbations on cellular metabolism and behaviors, since FBA simulates maximal efficiency of the metabolic system. We exemplify this method on two intestinal symbionts – *Bacteroides thetaiotaomicron* and *Methanobrevibacter smithii* – and visually consolidated the results into constellation diagrams that facilitate interpretation of information flow from given environments and thereby cultivate the design of controllable biological systems. The unique confluence of metabolic modeling and information theory in this model advances basic understanding of cellular metabolism and has applied value for the Internet of Bio-Nano Things, synthetic biology, microbial ecology, and autonomous laboratories.

## 1 Introduction

Engineering biological systems at the cellular-level is essential to realize the Internet of Bio-Nano Things (IoBNT) [1], where wearable bio-computers externally monitor and direct biological systems *in situ* [2]. These external perceptions utilize biosensing [3], optogenetics [4], or magnetic nanoparticles [5], but limitations such as off-target effects on the biological system and inherent biochemical noise compromise accuracy and liability of these technologies [6, 7]. The ideal method would instead leverage native metabolism to minimally disturb the studied system: e.g. ameliorating dysbiosis by changing the chyme nutrient flows via dietary intervention [8, 9]. The lack of basic knowledge of information flow, however, from an environment to behaviors in a metabolic network [10, 11] – post-translation modification, signal transduction [12], gene regulation, and biomass growth [6,10,13] – remains a bottleneck to realize this vision of environmentally controlling native metabolism [14].

Molecular communication (MC) theory in Figure S1 is an emerging confluence of communication and information theories [7, 15–17] that quantifies information flow (or mutual information) of an environment through black-boxed channels of chemical inputs to observed outputs [18]. The information flow *IF* [19] through each of these channels is quantified in binary digits, or bits (1 or 0),

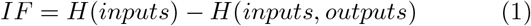

as the excess of input uncertainty *H*(*inputs*) from the output uncertainty *H*(*inputs, outputs*) from the given inputs. Flux Balance Analysis (FBA) in Figure S2 [20], by contrast, is a metabolic modeling framework that mechanistically predicts reactions fluxes and cellular behaviors of a metabolic system in a given environment [21] through linear optimization. FBA represents a metabolic system as a matrix *S* of stoichiometries for all metabolites in all reactions and determines the profile of reaction fluxes 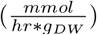 as a vector **v** that represents steady-state metabolic activity (*S***v** = 0) [22]. FBA optimizes for a sub-set of reaction fluxes (conventionally those contributing to biomass growth) and can be tailored through constraints on flux ranges (**v**_**min**_ ≤**v** ≤ **v**_**max**_) that can represent diverse chemical phenomena: e.g. reaction energetics [23] or regulatory processes [24].

MC and FBA methods offer complementary value for simulating the metabolic effects of an environment, however, these methods have never before been integrated into a single framework that quantifies information flow with mechanistic fortification. We therefore melded MC and FBA into a unique model that computes information flow through defined mechanically-resolved channels of metabolism. We defined a comprehensive MC channel from an input of consumed substrates to externally perceived behaviors such as excretions and biomass growth, and further partitioned this comprehensive channel into two intermediary channels [25]: one whose outputs are cytoplasmic reaction fluxes and one whose outputs are the externally perceived behaviors. These channels are illustrated in Figure S3. We exemplify our model on two [26] human symbionts that are associated with preventing metabolic diseases [27, 28]: the bacterium *B. theta* and the archaeon *M. smithii*. Seven substrates were selected for investigation on their propagation through the metabolism, which represent carbon, nitrogen, sulfur, and oxygen nutrient flows in chyme and have been previously studied with these organisms [26, 29]. The information flow of these substrates through these organisms was visualized in constellation diagrams [30] that elucidate the substrates that optimally transmits information through the metabolic networks. This model expands basic knowledge of cellular behavior as the causal consequence of environmental substrates, and can specifically address basic questions of molecular biology: 1) how much environmental information can a cell encode in its metabolism? and 2) how much information about intra-cellular metabolism can be externally perceived? These basic advances will accelerate biological engineering for myriad applications from healthcare to space colonization [31, 32].

## 2 Methods

Figure 1 compares each of the MC channels to the biological processes that they represent. The **Stage I** channel

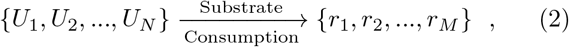

defines the activation of all *j* ∈ *M* cytoplasmic reactions *r*_*j*_ ∈ [0, 1] from the consumption of each substrate *U*_*i*_ ∈ [0, 1] for all *i ∈ N* compounds of interest, over a total of 2^*N*^ consumption profiles. The **Stage II** channel

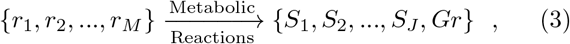

defines external excretions *S*_*e*_ ∈ [0, 1] for all *e* ∈ *E* exchangeable compounds and/or biomass growth *Gr* ∈ [0, 1] from the set of active reactions *r*_*j*_ ∈ [0, 1] ∀ *j* ∈ *M*. The **endto-end** channel is the entire pipeline that joins the **Stage I** and **Stage II** channels. The short-hand for each channel is therefore 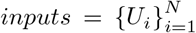 and 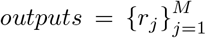 for **Stage I**; 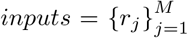 and 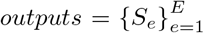, *Gr* for **Stage II**; and 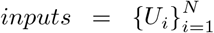 and 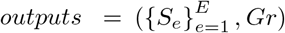 for **end-to-end**.

**Figure 1:**
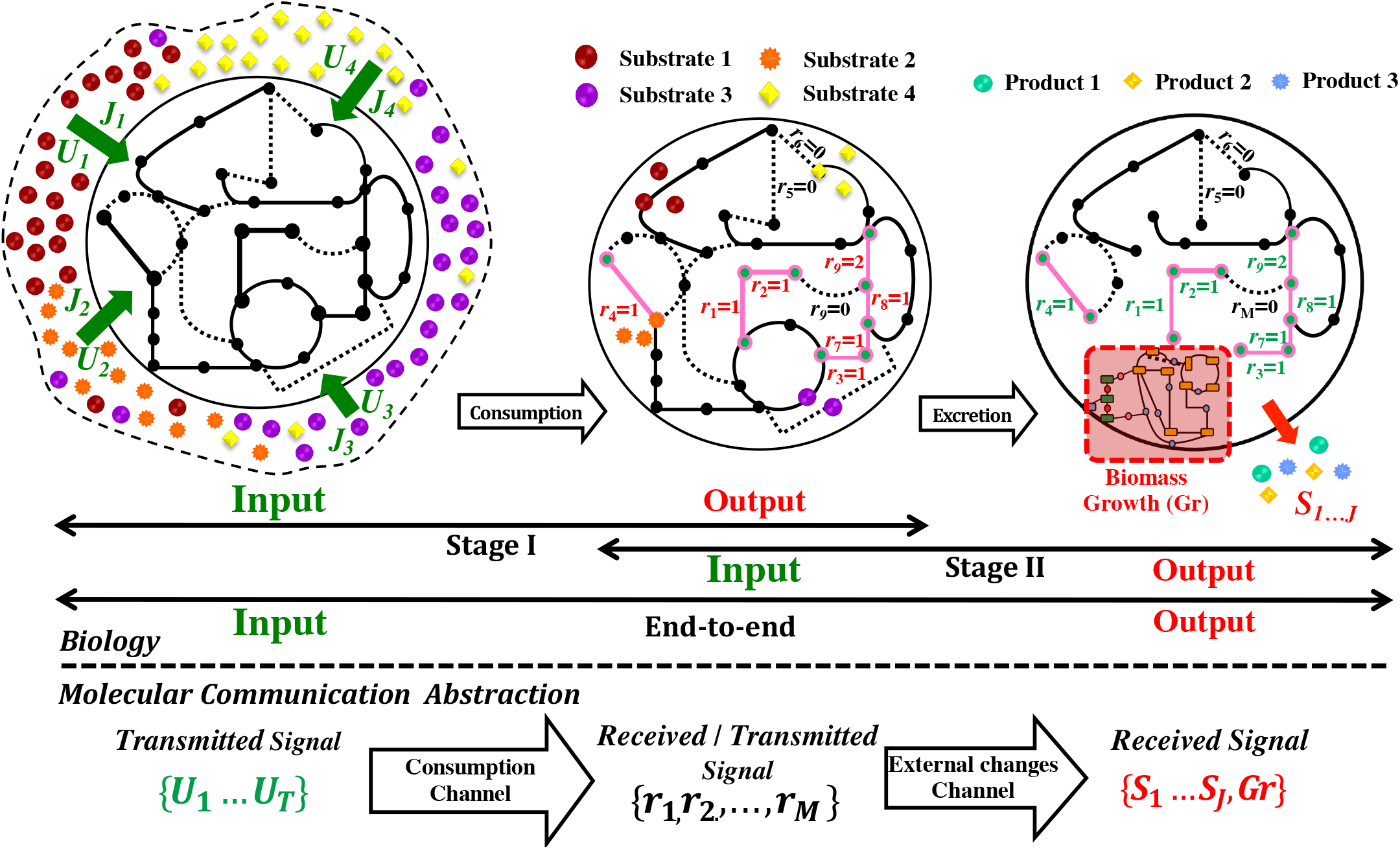
A comparison of biological metabolism and its molecular communication (MC) abstraction, above and below the dashed line, respectively. Biological pipeline depicts a) substrates consumption; b) the activation of metabolic pathways; and c) cellular outcomes such as excreta or biomass growth. The MC pipeline depicts these same phenomena as pairs of inputs and outputs through black-boxed channels: **Stage I** describes the activation of pathways from substrate consumption; **Stage II** describes the manifestation of cellular outcomes from the set of activated pathways; and the **end-to-end** channel describes the combination of **Stage I** and **Stage II** as the full pipeline.

### 2.1 Information flow

The input uncertainty *H*(*inputs*) from eq. (1) is generally defined as

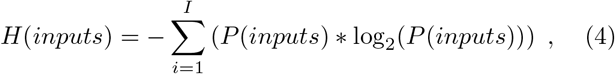

where *I* is the number of unique inputs and *P* () is a function that returns the probability of the provided argument, where we assume that all inputs and outputs are equally probable. The conditional output uncertainty *H*(*inputs, outputs*) is generally defined as

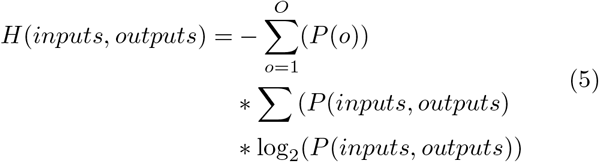

where *O* is the number of unique outputs. The *P* (*o*) term returns the probability of an output among all outputs, while *P* (*inputs, outputs*) returns the probability of an output per a given input. The summations in these equations become integrals when the inputs or outputs are continuous, respectively; however, our strict utilization of binary variables discretizes all domains and allows summations.

The presented general expressions for *H*(*inputs*) and *H*(*inputs, outputs*) are tailored for FBA simulations in several manners. First, the terms are demarcated with an asterisk 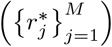 to denote that the value corresponds with the maximal information. Second, identical FBA solutions from different consumption profiles, based on their raw (continuous) fluxes, were grouped in Figure S4 to avoid redundant computations of *IF* ^*^. Concentration profiles were likewise grouped in Figure S5, where identical substrate profiles across multiple FBA simulations were consolidated.

### 2.2 Computational Tools and Workflow

The genome-scale models (GEMs) [33] of *B. theta* and *M. smithii*, which are detailed in Figure S6, were constructed through the KBase pipeline [34] in Figure 2. First, the organism’s genome sequences from the GenBank database [35, 36] were translated and mapped to functions via KEGG [37]. Second, these initial draft GEMs were gapfilled [38] in a media for each consumption profile (2^*N*^ differently gap-filled GEMs), where the minimal number of reactions that enables growth in the parameterized environment are added to the model. The seven examined compounds (*N* = 7) in our study, similar to other studies with these organisms [26, 29] – glucose (*G*), hematin (*He*), formate (*F*), *H*_2_, Vitamin (*B*_12_), acetate (*A*), and Vitamin K (*K*) – yielded 2^7^ = 128 consumption profiles for which *IF* ^*^ will be computed. The 128 unique consumption profiles were added to a standard base media that contained: *Ca*^+2^, *Mg*^+2^, *Cl*^−^, *Na*^+^, *K*^+^, *CO*_2_, *Co*^+2^, *Cu*^+^, *Fe*^+2^, *Fe*^+3^, *H*_2_*O, H*^+^, L-cysteine, *Mn*^+2^, *NH*^+^, *Ni*^+2^, *PO*^−^, that are common between the networks, such as Figure S8 of common metabolite-reaction connections between two FBA solution groupings.

**Figure 2:**
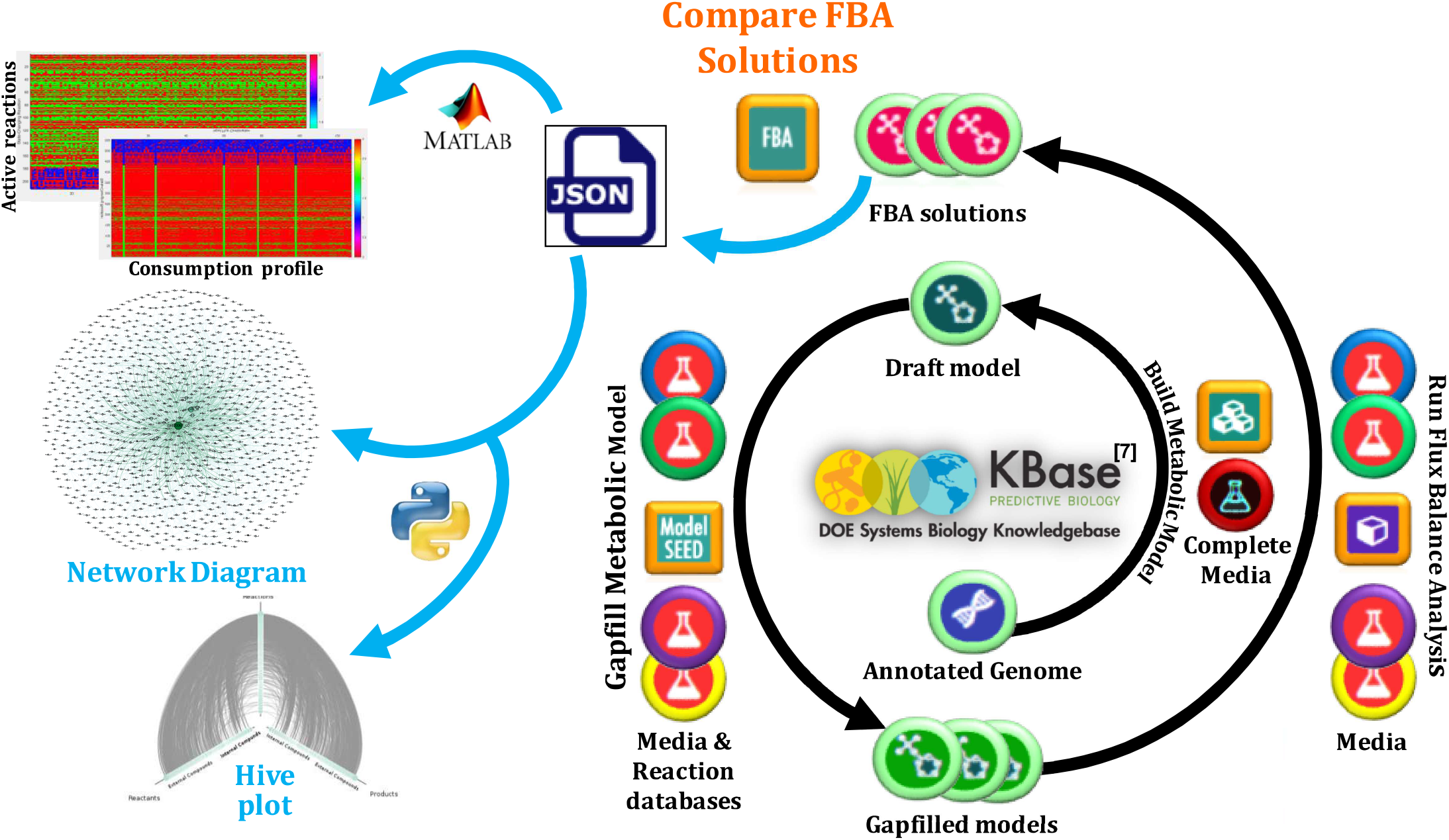
A summary of our modeling workflow. The initial steps leverage KBase as an open-source platform for conducting simulations and hosting our Application. Our KBase workflow involves: 1) annotating uploaded genomes through RAST or DRAMM; 2) reconstructing draft genome-scale metabolic models (GEMs) from the annotated genomes, via the `Build Metabolic Model` Application; 3) gap-filling the draft GEMs to create operational GEMs, via the `Gapfill Metabolic Model` Application; 4) acquiring FBA flux profiles in the parameterized media, via the `Run Flux Balance Analysis` Application; and then 5) exporting a JSON that compares FBA solutions of the different GEMs and conditions via the `Compare FBA Solutions` Application. Custom MATLAB and Python scripts then process the exported JSON of compared FBA solutions into heatmaps that illustrate activated pathways and into network diagrams and hive plots to illustrate input-output relationships in the metabolism, respectively.

## 3 Numerical Results

The maximal information flow *IF*^*^ for each channel was determined by computing each term in eq. (1) according to the Stage channels that are defined in eqs. (2) and (3). The as-sumed equal probability of the *N* = 7 examined substrates is 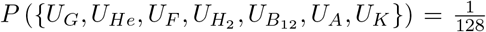. The input uncertainty in eq. (4) for Stage I and the 128 possible inputs was computed: *H*(*inputs*) = 1 log_2_(128) = 7 *bits*. The conditional output uncertainty for Stage I of *B. theta* was calculated, from 113 unique reaction activation profiles (*M* = 113) over 14 unique FBA solutions,

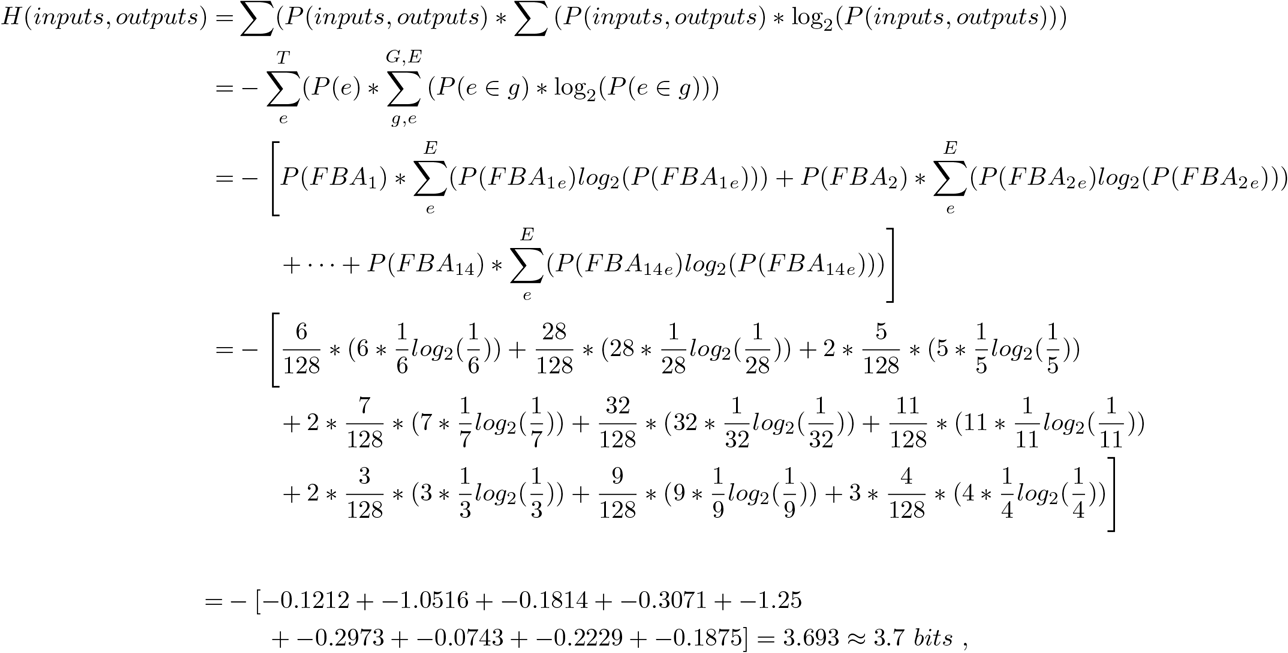

where the uncertainty of each FBA group is separately calculated and then aggregated into the total output uncertainty. The grouping of the 128 FBA solutions based on reaction fluxes in Figure S4 importantly simplifies the above computation and can facilitate biological investigation in relating consumed substrates to metabolic activation and cellular behaviors. The equivalent probability of a FBA group consumption profile (element *e*) among all other consumption profiles in the total set of consumption profiles (*T*) or in the same FBA group 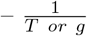– allows the terms for all elements to be condensed with a coefficient of the number of elements in the group: e.g. the total uncertainty for the 6 elements of FBA group 1 (*FBA*_1_) is 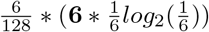, where the coefficient **6** collapses the six element terms into a single term. Several FBA groups also possessed the same number of elements, which permitted further simplification with a coefficient of the number of identically sized FBA groups, where all FBA groups are themselves equally probable: e.g. FBA groups 3 and 4 each possess five elements and can therefore be jointly defined as 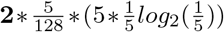, with the coefficient **2** denoting the two FBA groups.

The *IF*^*^ for Stage I of *B. theta* then finally equates *H*(*inputs*) − *H*(*inputs, outputs*) = 7 − 3.7 = 3.3 *bits*. The conditional output uncertainty for Stage I of *M. smithii* was computed to be 2.5 *bits*, analogously to the above derivation for *B. theta* except with 136 unique reaction activation profiles (*M* = 136) over 31 unique FBA solutions. The *IF*^*^ for Stage I of *M. smithii* then equates 7 − 2.5 = 4.5 *bits*. These results are depicted Figure S9, where *B. theta* reaches its maximum information flow in the absence of formate while *M. smithii* reaches its maximum information flow with all seven examined substrates.

The FBA groupings that were defined for the Stage I calculations served as the inputs for Stage II and were used to compute the input uncertainty: −1 log_2_(14) = 3.8 *bits* for *B. theta* and −1 * log_2_(31) = 5.0 *bits* for *M. smithii*. The FBA solutions were then regrouped according to the respective output – *Gr* (*outputs* = *Gr*) or both *S*_*e*_ ∀ *e* ∈ *E* and *Gr* (*outputs* = (*S*_*e*_ ∀ *e ∈ E, Gr*)) – to calculate the conditional output uncertainty of the Stage II channel. Grouping by biomass growth (*outputs* = *Gr*) in Figure 3a separately evaluated the probability of each group *g* in the set of all FBA groups *G*. The Stage II *IF*^*^ for biomass growth is therefore 3.8 − = 2.8 *bits*, and analogously *IF*^*^ = 5 − 1.1 = 3.9 *bits* for *M. smithii*. Grouping by both excretions and biomass growth manifested in *IF*^*^ = 3.8 −.1 = 3.7 for *B. theta* and *IF*^*^ = 5.0 − 1.1 = 3.9 for *M. smithii*. The modest increase in *IF*^*^ for *B. theta* when including metabolic excretion to the outputs, and the unchanged *IF*^*^ of *M. smithii*, is counter-intuitive, since more outcomes would seem to capture more inputs; evidently, biomass growth captures most of the information flow during substrate consumption through metabolism.

**Figure 3:**
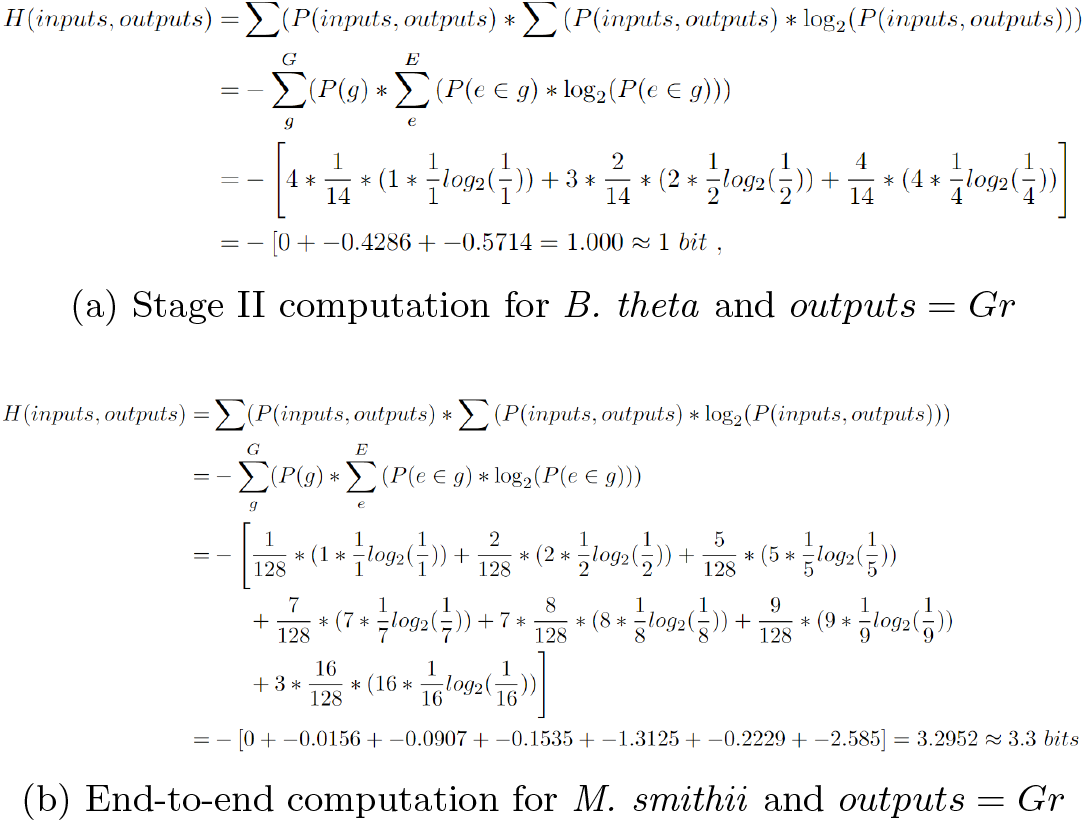
*IF*^*^ computations for Stage II and end-to-end channels for the two examined organisms.

The end-to-end channel – *inputs* = 128 *consumption profiles, outputs* = *Gr*, and FBA groups according to *Gr* – was additionally computed to be *IF*^*^ = 7 − 4.3 = 2.7 *bits* for *B. theta* and *IF*^*^ = 7− 3.3 = 3.7 *bits* for *M. smithii* in Figure S10, where the conditional output was computed in fig. 3b for *M. smithii*. The subtly lower *IF*^*^ values are expected, since the larger scope of this channel introduces more noise and loss that hinder reception of the signal in the final outputs.

### 3.1 Visualizations

The constellation diagrams of Figure 5 statistically compress the *IF*^*^ from all consumption profiles for rapid interpretation. These diagrams were constructed by first: 1) grouping consumption profiles by *IF*^*^ from the end-to-end channel; 2) determining the minimum and maximum *IF*^*^ for each consumption profile; and 3) graphing each of the minimum, maximum, and average *IF*^*^ (excluding the minimum and maximum) by the size of the consumption profile. These diagrams interestingly revealed that the maximum *IF*^*^ plateaus with ≥ 5 of the examined substrates, which suggests that there are diminishing returns to cellular control from environmental perturbations. This trend is consistent with simulations from the BioSIMP software that identified substrates for growth of *B. theta* and *M. smithii* [29]. This trend is moreover echoed by Figures 4, S9, and S10 that reveal relatively small consumption profiles with remarkably high *IF*^*^: e.g. consuming *F, B*_12_, and *A* for *B. theta* and *M. smithii* ; or consuming *F, H*_2_, *A* for *M. smithii*. These basic insights from various substrate conditions would be necessary for biomedical monitors and other systems that use molecular transport to categorize metabolic states.

**Figure 4:**
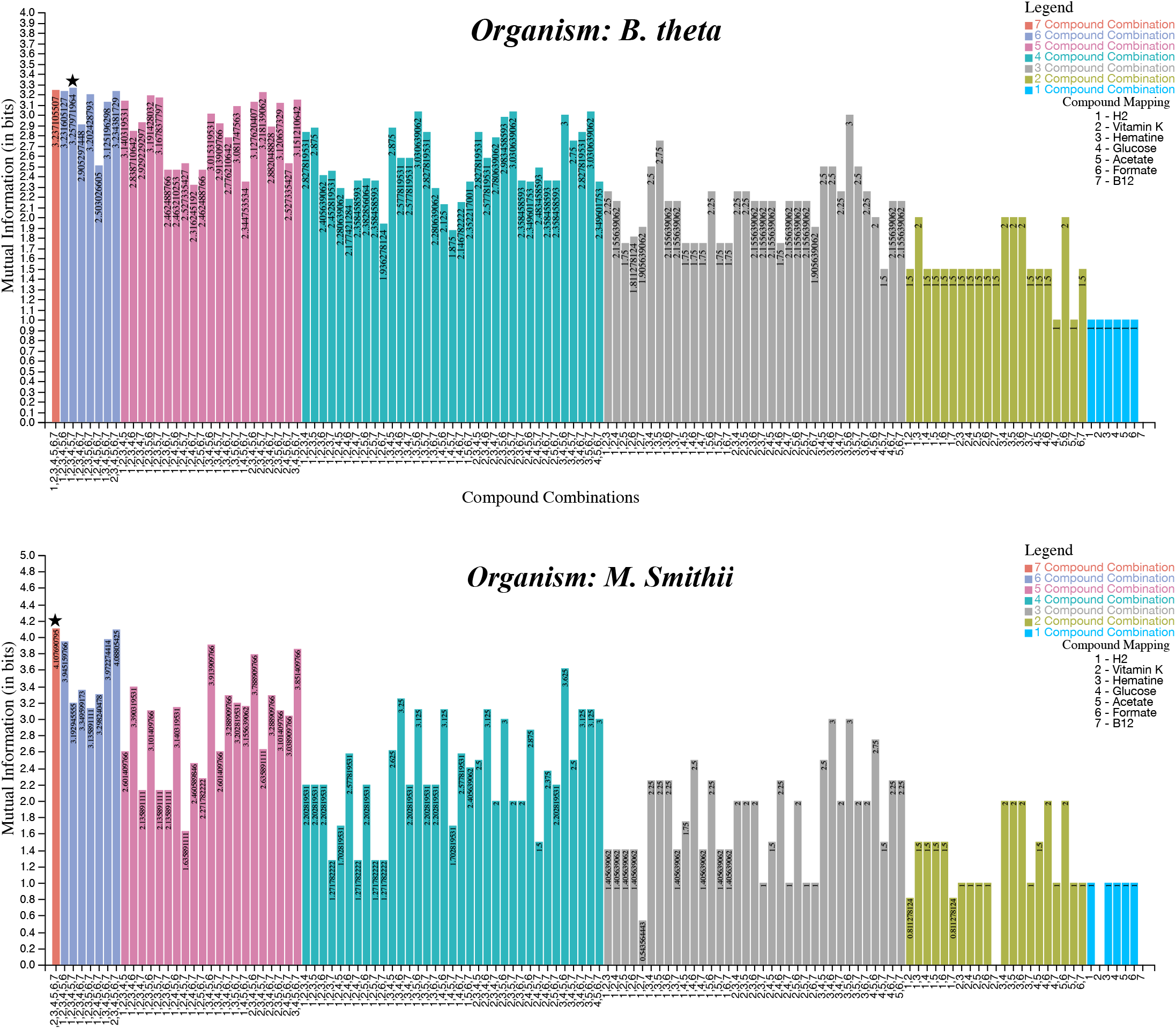
The end-to-end upper bounds for each consumption profile. The maximal information flow is starred. The consumption profiles with more compounds predictably manifested in greater potential information flow. The consumption profiles are condensed on the x-axis by numbering each substrate: 1 = glucose (*G*), 2 = hematin (*He*), 3 = formate (*F*), 4 = *H*_2_, 5 = Vitamin (*B*_12_), 6 = acetate (*A*), and 7 = Vitamin K (*K*).

**Figure 5:**
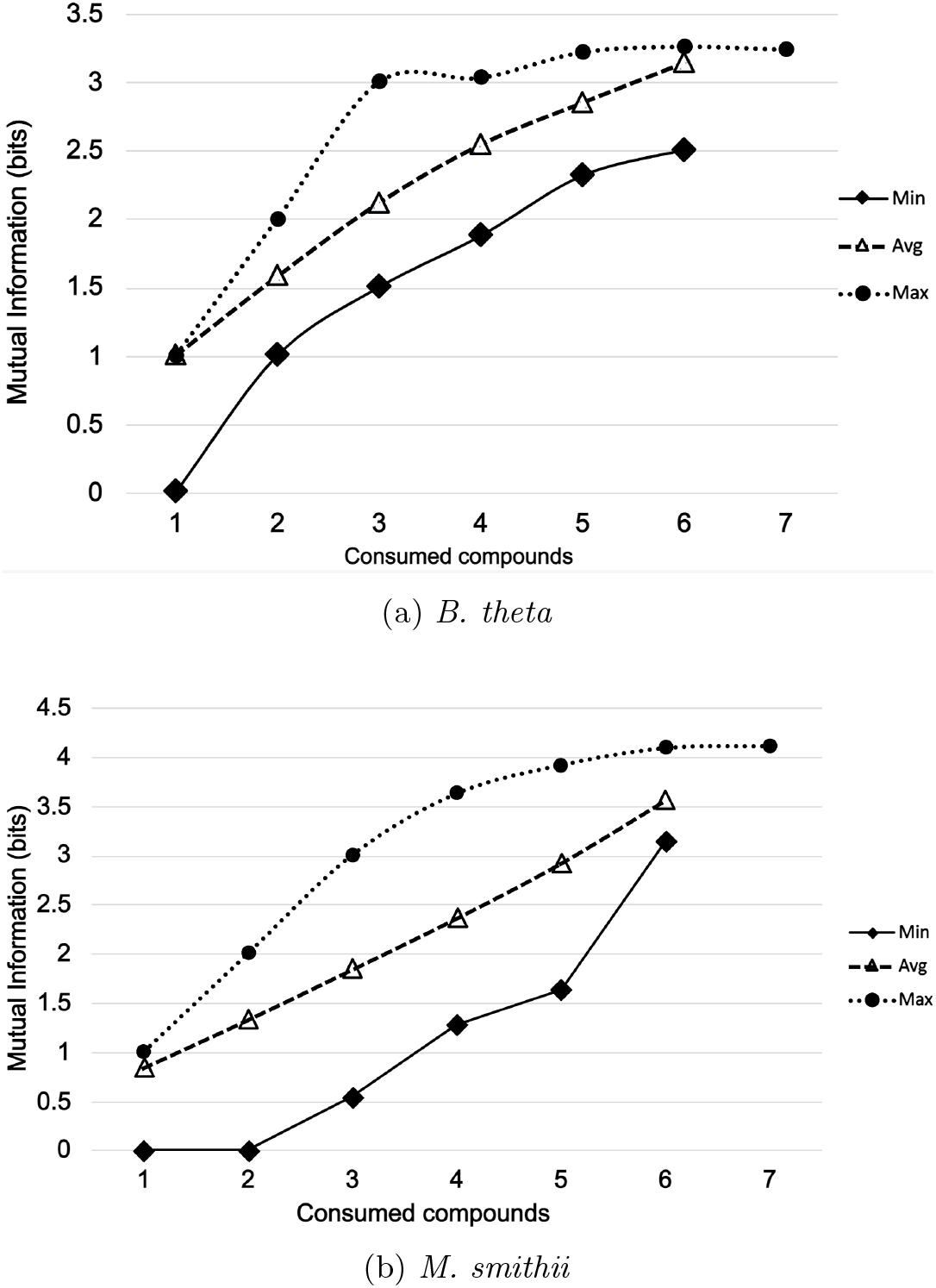
Constellation diagrams for the information flow from all consumption profiles in the end-to-end channel. The minimum, maximum, and average values are plotted against the number of compounds in the consumption profiles, which reveals that the maximum *IF*^*^ increases logarithmically while the average *IF*^*^ increases linearly. This observation is intuitive where biochemical noise over the channel defines a fundamental limit to *IF*^*^ but generally an increase in the degrees of freedom (consumed substrates) increases the uncertainty (*IF*^*^) of a system.

## 4 Conclusions and Future Work

Molecular communication theory quantifies the flow of information from extracellular environments into metabolic activity, substrate excretions, or biomass growth. We merge this concept with mechanistic FBA, and defined several metabolic channels through which environmental information likely flows in a cell, to compose a unique model for quantifying optimal information flow through metabolic systems. Our model and constellation diagrams of the results identified consumption profiles that optimally transmitted information biochemically to cellular outcomes, such as biomass growth and chemical excretions, which can minimize the number of resource-intensive experiments that are needed to find substrate profiles that desirably control cellular behaviors. Hive plot visualizations of our results further conveyed connectivity between inputs and outputs, which empowers more detailed analyses. The Code, KBase Narratives, and additional details are available in our GitHub repository (https://github.com/freiburgermsu/MetabolicMC_Supplementary_Git_v1).

This model advances basic biological knowledge of cellular engagement with its environment, and provides a tool to identify environmental substrate combinations that can most effectively engineering and monitor cellular behavior. This tool has diverse applications in biomedical wearable technologies and ecological control through benign substrate perturbations. Subsequent work will refine our model by (1) better compensating biochemical noise; 2) capturing dynamic information flows and continuous variables via the dynamic FBA algorithm; 3) validating predictions with experimental data; and 4) exploring applications in autonomous laboratories that predictably monitor and control cellular behaviors through environmental perturbations.

## Acknowledgment

This work was funded by the US National Science Foundation through grant MCB-1449014, the National Institutes of Health 11-P20-GM113126-01, hi Sa the NSF EPSCoR EPS1004094 and the CCF-1161767. Also the grant supported by NIH 11-P20-GM113126-01. The authors would also like to thank KBase developers for their active support throughout the development of this research.

## Supplementary Information

**Figure S1:**
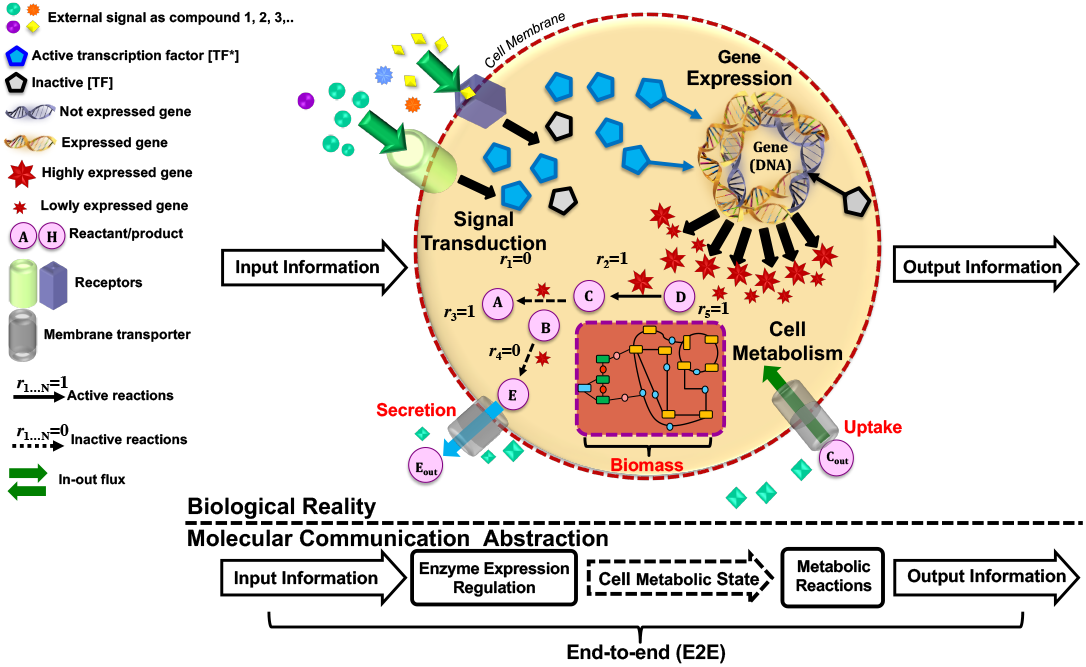
Biological reality of Molecular communication in cell metabolism.

**Figure S2:**
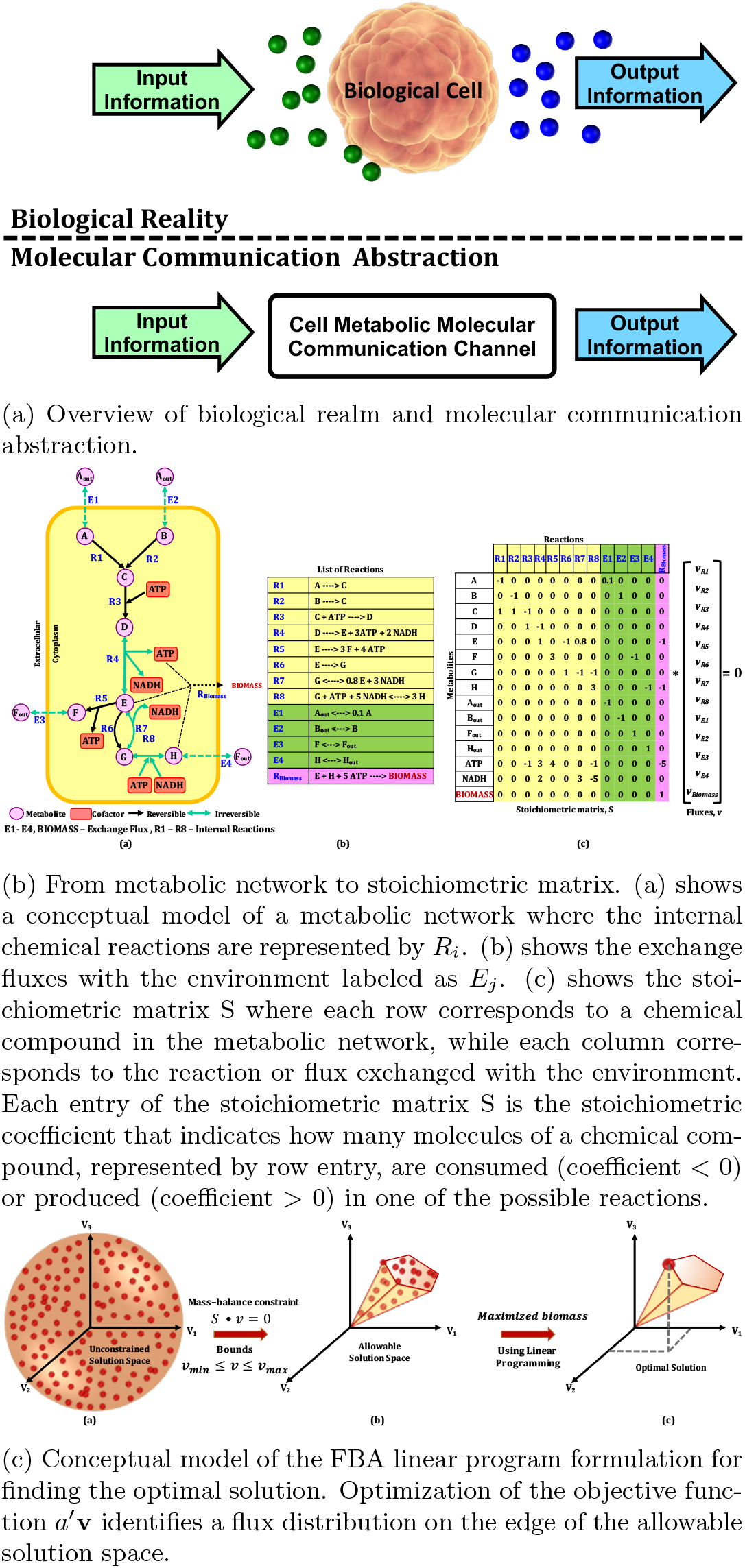
Background information on a) the molecular communication abstraction of cellular systems, where cellular metabolism is black-boxed; b) the FBA algorithm and its use of matrix algebra; and c) the geometric solution space that linear optimization in FBA navigates to find the optimal solution.

**Figure S3:**
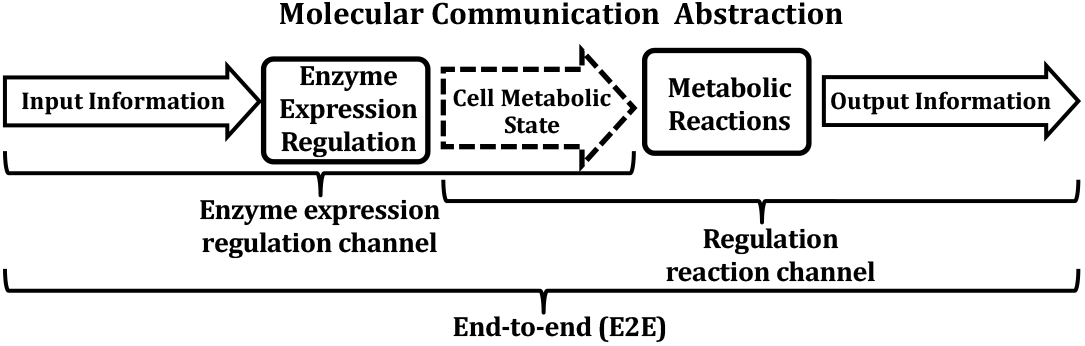
Molecular communication channel abstraction of cell metabolism.

**Figure S4:**
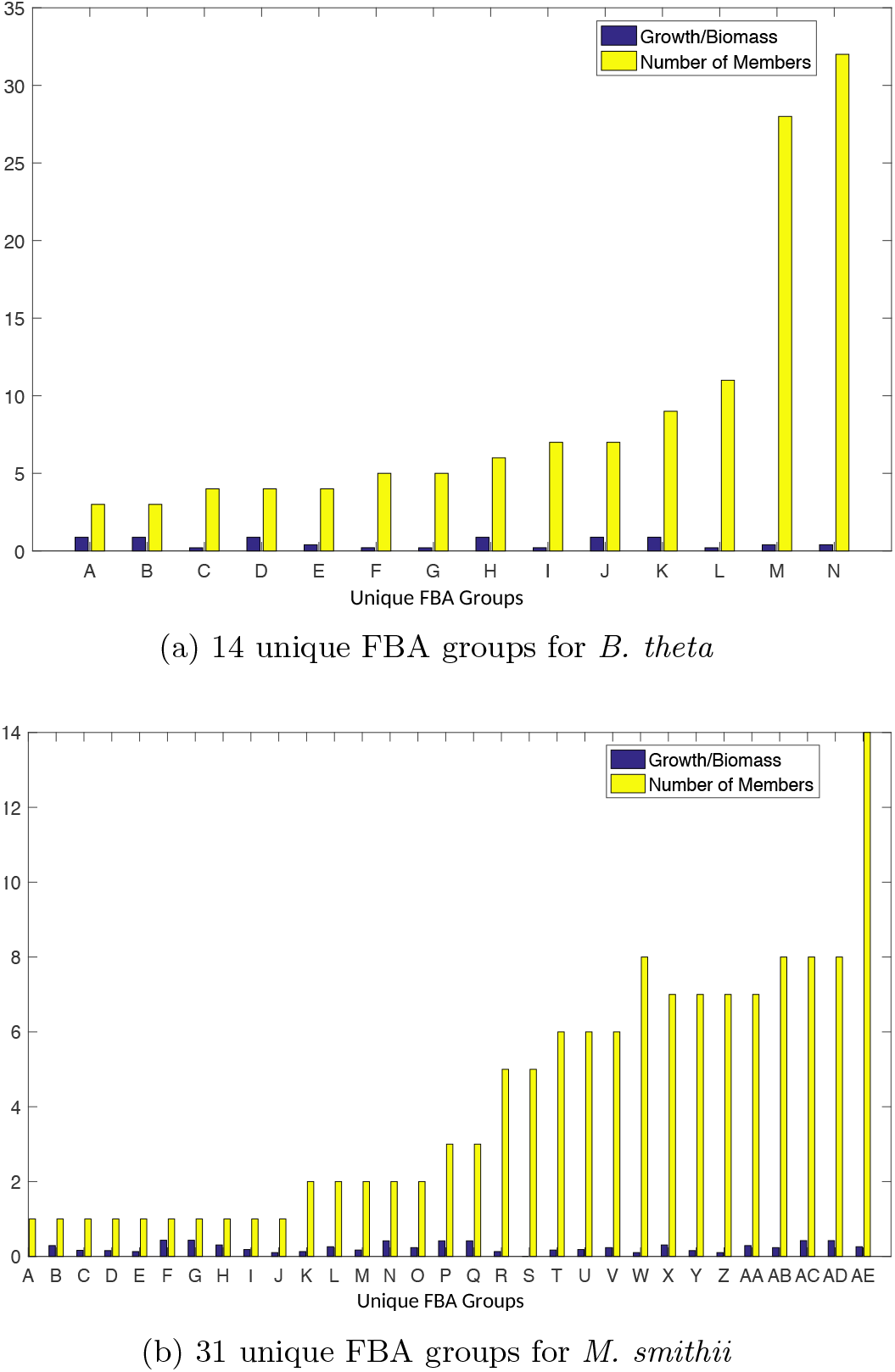
The number of FBA solutions per group of unique activation profiles, over all consumption profiles. The *M. smithii* metabolism exhibited more than twice as many unique metabolic responses to the consumption profiles than *B. theta*, which suggests that it is more dynamic to its environment. This is corroborated by the much higher proportion of active reactions in Figure S5 relative to *B. theta*.

**Figure S5:**
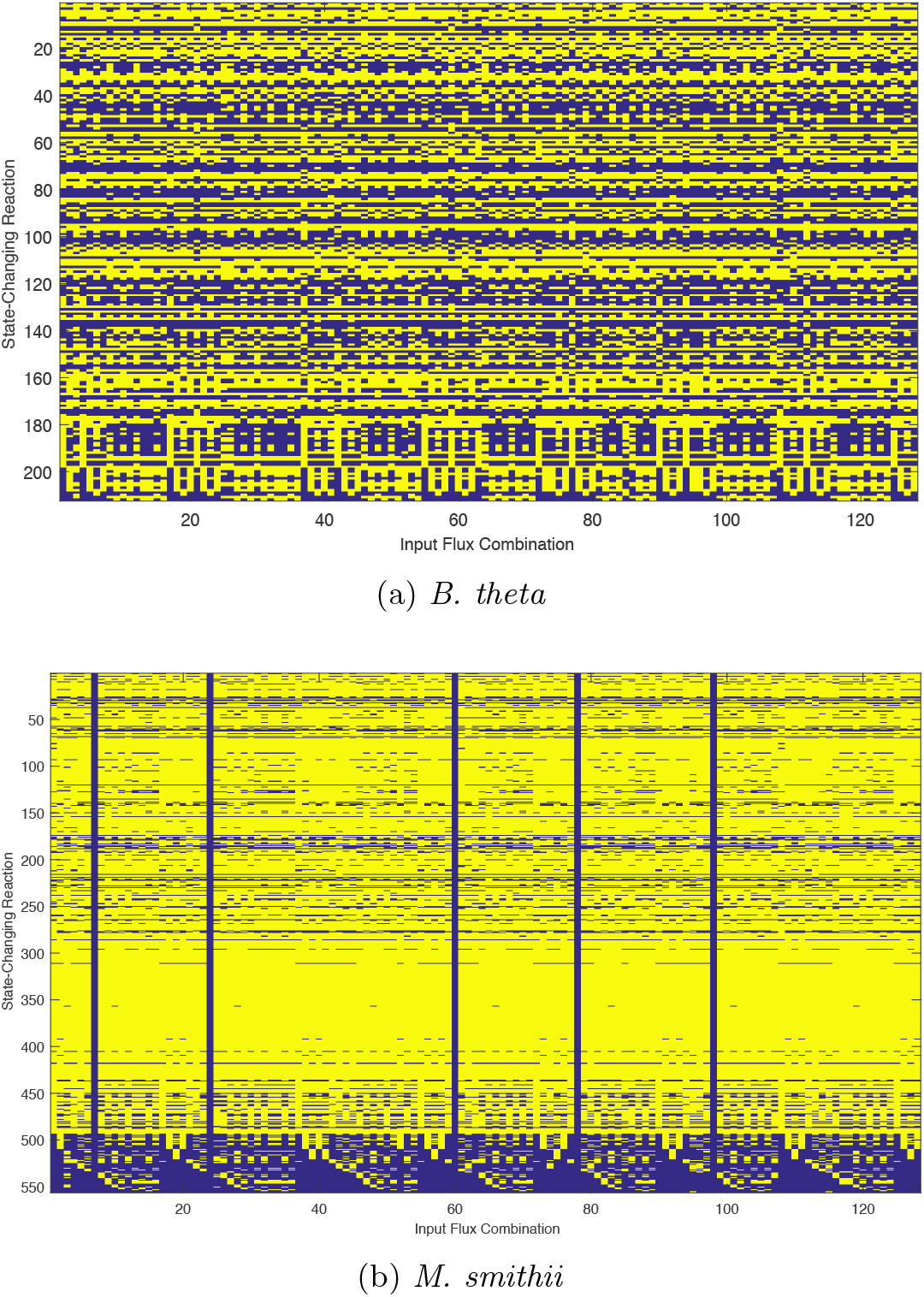
Reaction activation from FBA simulations 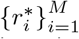 for each possible consumption profile, where yellow is active and violet is inactive. *M. smithii* metabolism is evidently much more activated than *B. theta* by the examined media, notwithstanding a handful of media profiles that did not support growth (illustrated as vertical lines), although *B. theta* seems to have exhibited at least some metabolic activation in all of the examined media. This may be the consequence of *B. theta* utilizing anaerobic fermentation that expands its habitable zone beyond that of *M. smithii*.

**Figure S6:**
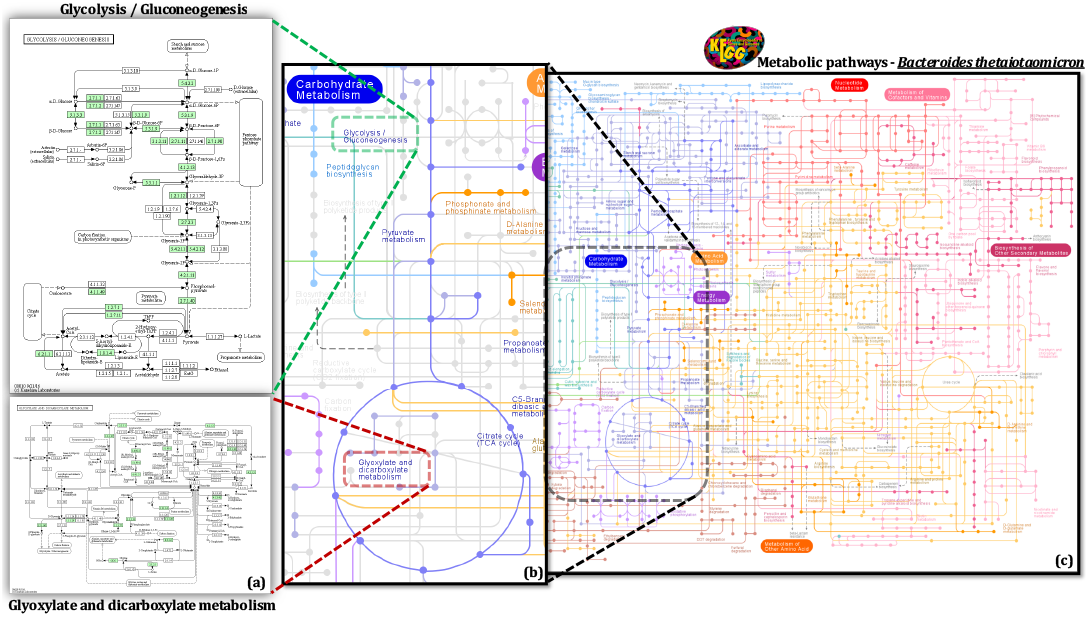
KEGG module of *Bacteroides thetaiotaomicron* metabolic pathways. **(a)** Summary of the biological processes shown in the pathway map of Glycolysis / Gluconeogenesis and Glyoxylate and dicarboxylate metabolism. **(b)** Enlarged fine details of a section of a complete metabolic model. **(c)** Part of the complete KEGG database pathway maps of *Bacteroides thetaiotaomicron*. visualized parts of a GEM (on the right) for the organisms *B. theta*, used in our study, which is obtained from the Kyoto Encyclopedia of Genes and Genomes (KEGG) database [37]. The nodes represent compounds that are inputs/outputs to the reactions, and edges represent the chemical reactions. Inputs from the environment taken by the organism are involved in the reactions of metabolic pathways, resulting in the exchange of fluxes with the environment (uptake and secretion) or in the production of biomass (growth).

**Figure S7:**
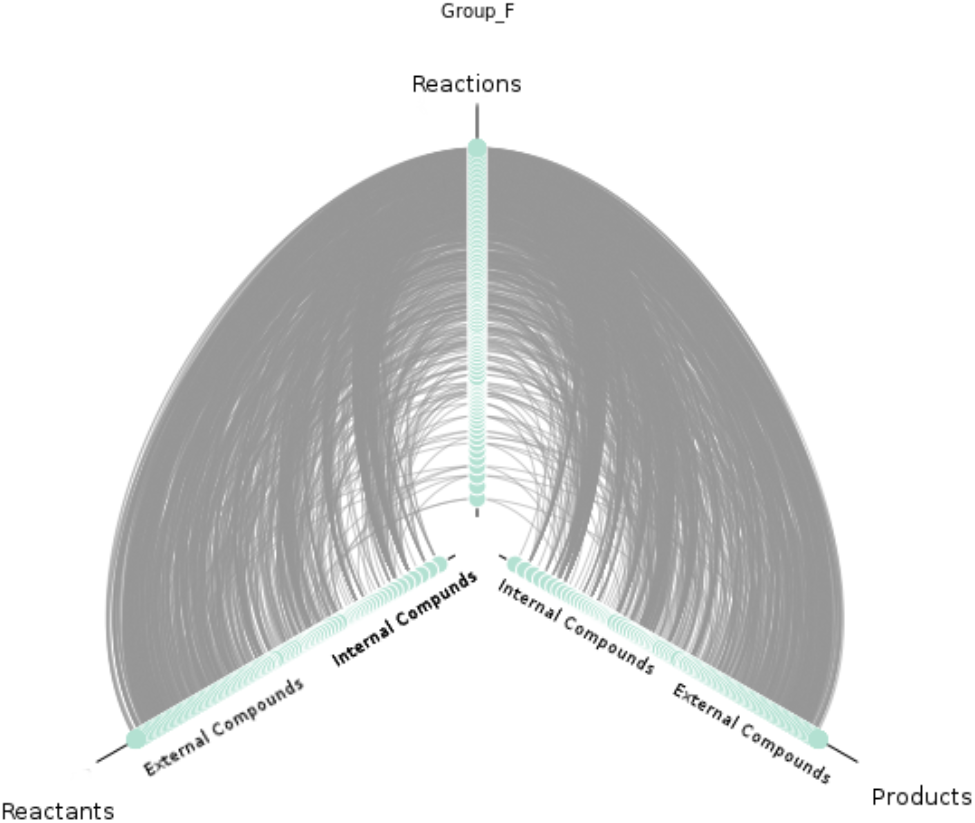
A hive plot for a configuration F is shown in the figure. The reactions are placed on the Z axis, the reactants on the X axis and the products on the Y axis. Further the External compounds are placed higher on the X and Y axes than the Internal compounds.

**Table S1:** **v**_**max**_ of minimal media compounds and important 7 compounds.

| Compound Name | MaxFlux (mmol/g CDW/hr) |
| --- | --- |
| Calcium ($\text{Ca}^{2+}$) | 0.180254 |
| Chloride ($\text{Cl}^-$) | 16.058 |
| Carbon dioxide ($\text{CO}_2$) | 34.00204 |
| Cobalt ($\text{Co}^{2+}$) | 0.042029 |
| Copper ($\text{Cu}^{2+}$) | 1 |
| Ferrous ($\text{Fe}^{2+}$) | 0.014 |
| $\text{H}^+$ | 1 |
| Water ($\text{H}_2\text{O}$) | 46166.89 |
| Potassium ($\text{K}^+$) | 100 |
| L – Cysteine | 2.8 |
| Magnesium (Mg) | 0.098375 |
| Manganese ($\text{Mn}^{2+}$) | 0.050529 |
| Sodium ($\text{Na}^+$) | 17.564 |
| Ammonium ($\text{NH}_3$) | 7.5 |
| Nickel ($\text{Ni}^{2+}$) | 1 |
| Phosphate | 100 |
| Sodium bicarbonate | 11.9 |
| Sulfate | 7.5 |
| Zinc ($\text{Zn}^{2+}$) | 1 |
| Ferric ($\text{Fe}^{3+}$) | 0.014 |
| D-Glucose | 2.78 |
| Hematin | 0.02 |
| Formate | 10 |
| $\text{H}_2$ | 0.78000078 |
| Vitamin B12 | 0.0000037 |
| Acetate | 10 |
| Vitamin K | 5.8 |

**Figure S8:**
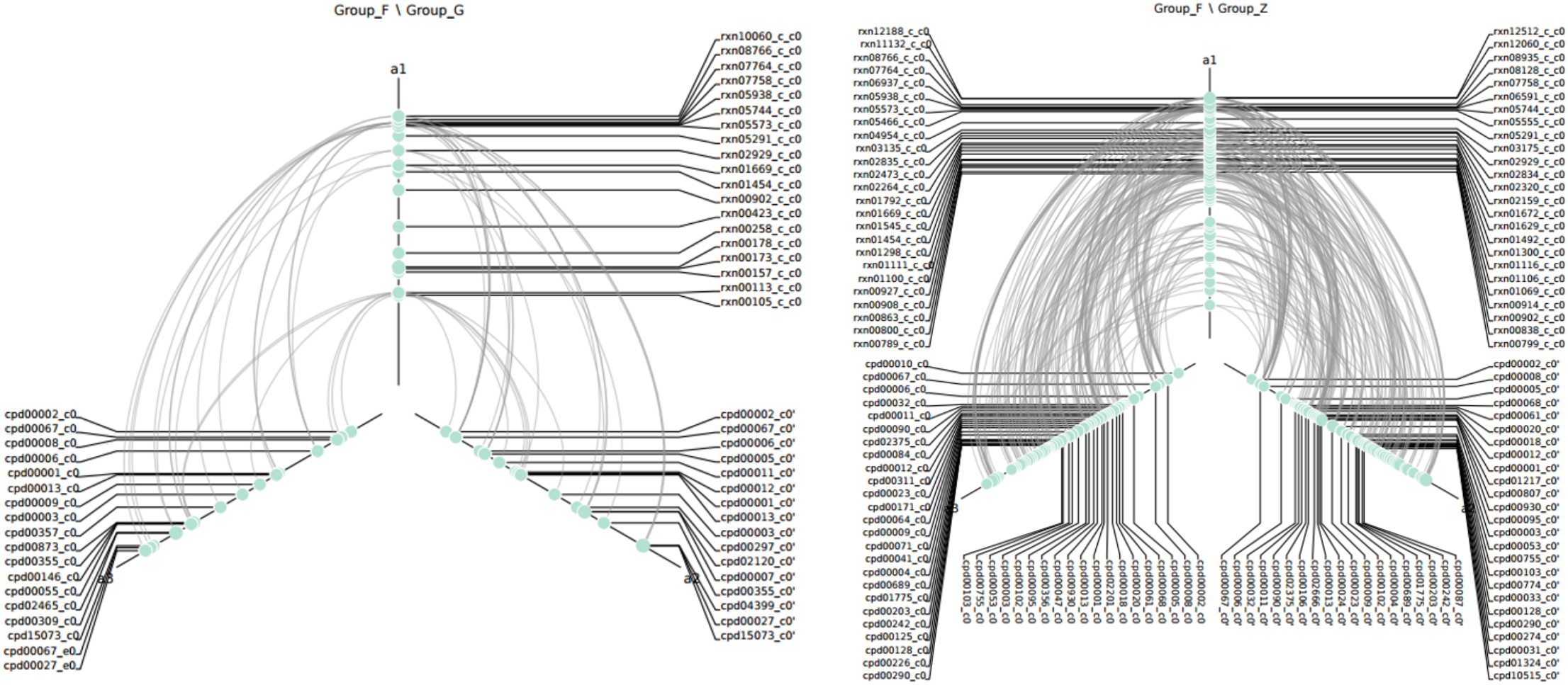
In this figure we take 3 different configuration of state changing reactions, labelled F, G and Z. It shows differential hive plots of F vs G and F vs Z. The groups F and G in F vs G hive plot has the same biomass whereas, the groups F and Z in F vz Z hive plot have the least and highest biomass respectively. When a reaction is present in F and absent in G or Z the reaction is represented along with its links to the compounds. When a reaction is present in the other groups but absent in group F the reaction is shown as a node not connected to any other compounds.

**Figure S9:**
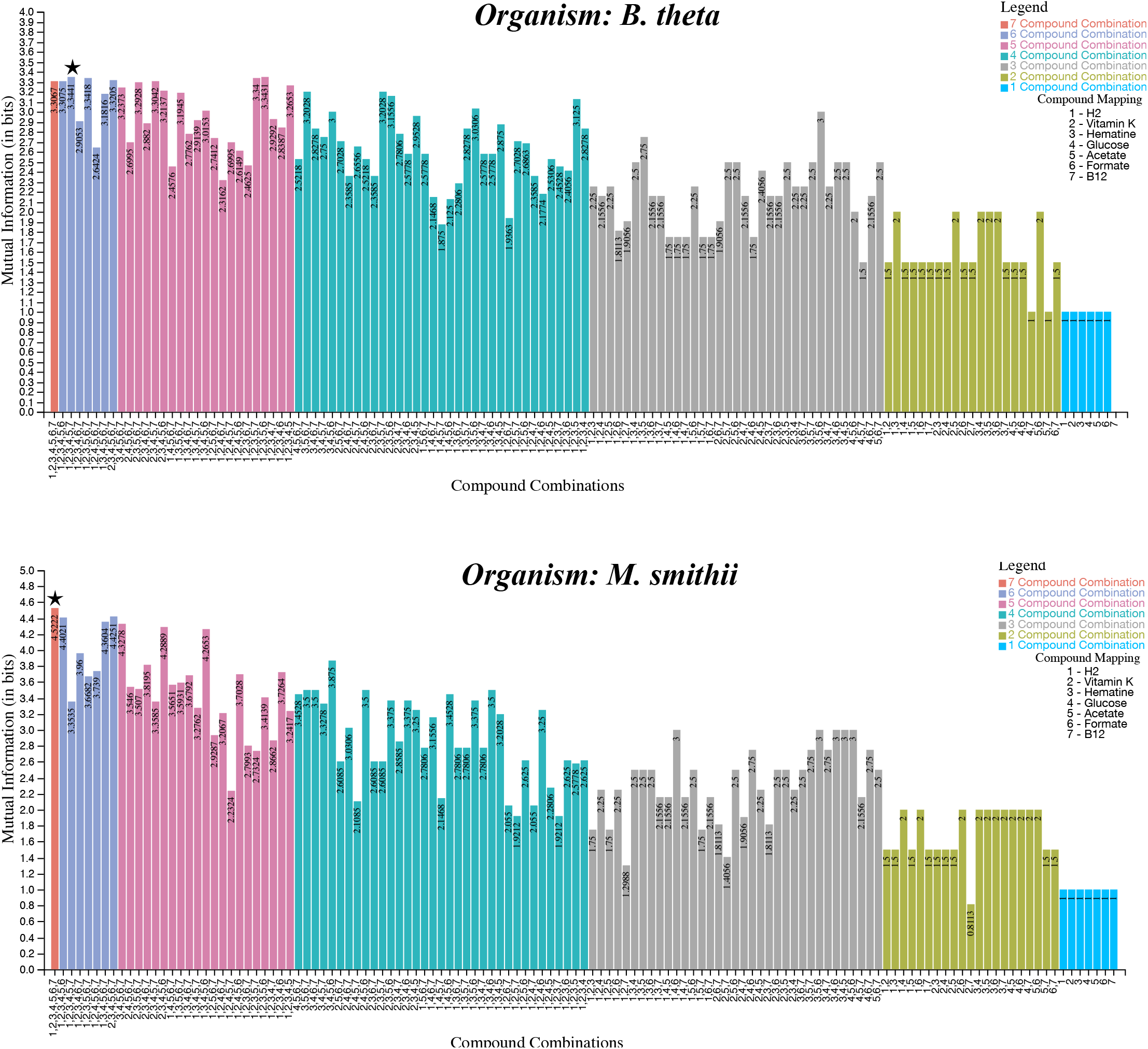
Upper bounds of the steady-state mutual information for all the different combinations of seven compounds in Enzyme Expression Regulation channel in. The best estimation of the maximum information is starred.

**Figure S10:**
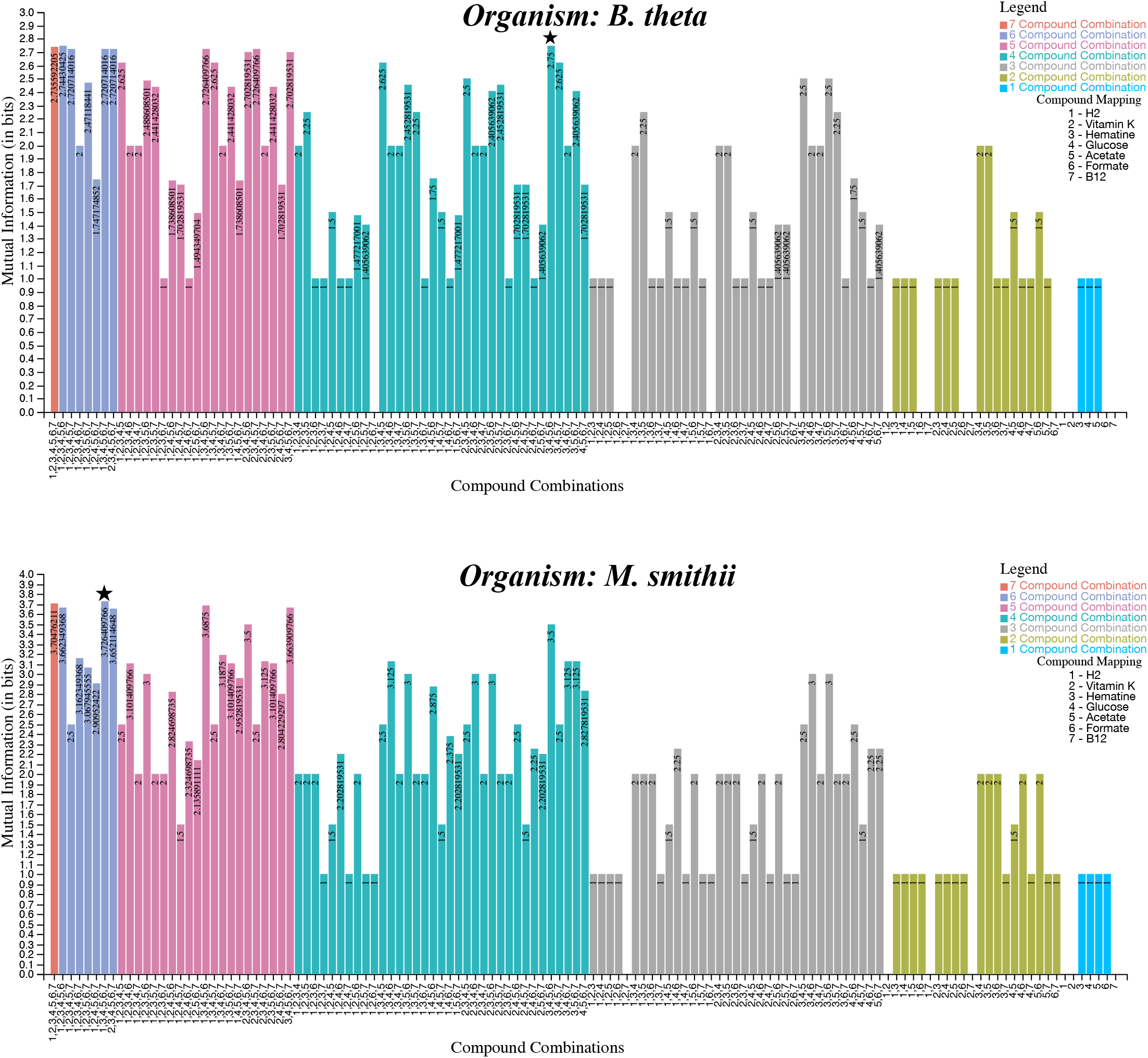
Upper bounds of the steady-state mutual information for all the different combinations of seven compounds in E2E in with respect to **Biomass only**. The best estimation of the maximum information is starred.

**Figure S11:**
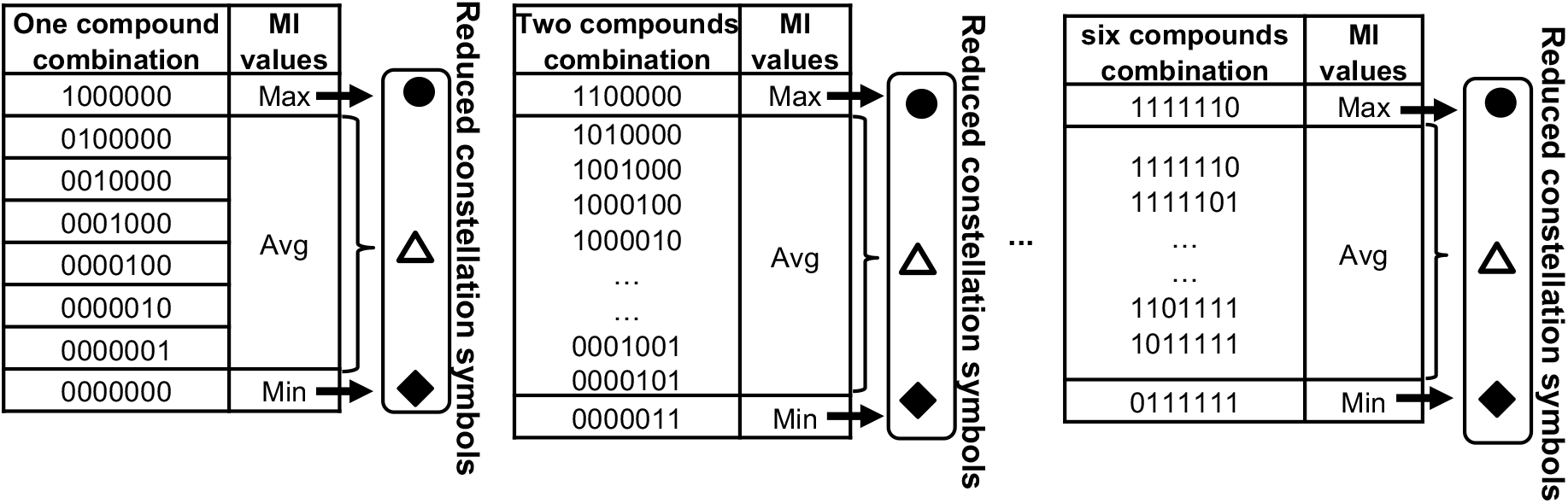
Steps involved in constructing constellation diagram.

